# A Humanized IFN-γ Mouse Model Reveals Skin Eschar Formation, Enhanced Susceptibility and Scrub Typhus Pathogenesis

**DOI:** 10.1101/2025.08.01.668084

**Authors:** Ryan H. Cho, Lihai Gao, Casey Gonzales, Dario Villacreses, Emmett A. Dews, Hema P. Narra, Hui Wang, Lynn Soong, Yuejin Liang

**Affiliations:** Department of Microbiology and Immunology, University of Texas Medical Branch, Galveston, Texas, USA; Department of Pathology, University of Texas Medical Branch, Galveston, Texas, USA; Institute for Human Infections and Immunity, University of Texas Medical Branch, Galveston, Texas, USA; Center for Tropical Diseases, University of Texas Medical Branch, Galveston, Texas, USA

**Keywords:** *Orientia*, interferon, eschar formation, intradermal inoculation, humanized mice

## Abstract

Scrub typhus, caused by *Orientia tsutsugamushi* (*Ot*) bacteria, is a serious acute febrile illness associated with significant mortality. An estimated one million cases occur annually, with more than one billion people at risk. No effective vaccine is currently available, largely due to the complex *Ot* strain diversity and an incomplete understanding of protective immune mechanisms. To overcome these challenges, there is a critical need for a suitable animal model that mimics human disease through the natural route of infection via mites. Here, we report for the first time that a genetically engineered humanized mouse strain (with triple knockout/knock-in of IFN-γ and its receptors), exhibits increased susceptibility to intradermal *Ot* infection compared to wild-type (WT) mice. This is evidenced by greater body weight loss, elevated bacterial burden, and reduced expression of interferon-stimulated genes (ISGs). Humanized mice exhibit pronounced biochemical abnormalities and tissue pathology accompanied by dysregulated T cell and neutrophil responses following infection. Notably, these immunocompetent mice develop skin eschar-like lesions resembling those observed in human patients. Overall, our study introduces a promising humanized mouse model to dissect the immunopathogenesis of scrub typhus and evaluate future vaccine candidates.

**Author Summary:** Scrub typhus is a serious disease caused by the obligately intracellular bacterium *Ot* that spreads to humans through the bite of larval mites called chiggers. It affects over a million people each year, primarily in Asia, and can lead to life-threatening complications. Unfortunately, we still lack a clear understanding of how this infection causes disease, partly because there is not a good laboratory model that closely reflects how humans respond to infection. Our recent reports have suggested an important role of IFN-γ in host protection against *Ot* infection. In this study, we used a new genetically modified mouse strain that carries human IFN-γ signaling in place of its mouse counterpart. We found that these humanized mice are more vulnerable to infection, develop skin eschar lesions like those in patients, and show signs of systemic inflammation and organ damage. Their immune response also resembled what has been observed in human patients. This new mouse model can help scientists better understand the mechanisms as to how this bacterial species causes severe disease outcomes in patients. Once those mechanisms are understood, this mouse model will serve a further purpose as a tool for testing new vaccines and treatments to aid humans at-risk for scrub typhus.

## Introduction

*Orientia tsutsugamushi* (*Ot*) is an obligately intracellular bacterium that causes scrub typhus in humans. An estimated one million cases occur annually, with over one billion people at risk, mostly within the “tsutsugamushi triangle”, which extends from Pakistan in the west to far eastern Russia in the east to northern Australia in the south(1, 2). *Ot* is primarily transmitted to humans through the bite of infected chigger mites of the *Leptotrombidium* genus, such as *Leptrotrombidium palpale,* a reservoir which maintains the bacterium through transovarial transmission(3). The clinical manifestations of scrub typhus include fever, headache, myalgia, vomiting, lymphadenopathy, and the characteristic eschar at the site of the chigger bite(4–6). In severe cases, bacterial dissemination to multiple organs can result in life-threatening complications such as respiratory failure, disseminated intravascular coagulation, meningitis, and encephalitis(7–9). Although traditionally endemic to countries within the tsutsugamushi triangle, scrub typhus cases have been reported in Africa, South America, and the Middle East(10). Notably, a recent discovery of *Orientia* DNA in chiggers in North Carolina raises concerns about potential emergence in the United States(11). Currently, there is no licensed vaccine for scrub typhus, partly due to a limited understanding of host-pathogen interactions and the extensive antigenic diversity among *Orientia* strains(12, 13). This lack of immunological insight leaves us ill-prepared to combat this neglected mite-borne disease, underscoring the urgent need to elucidate the mechanisms underlying *Ot* pathogenesis.

Animal models that closely recapitulate human scrub typhus are essential for advancing immunological studies and evaluating vaccine candidates. Reported animal models include intraperitoneal, intravenous, and intradermal routes of infection in mice, as well as intradermal and live chigger challenge models in non-human primates(14–22). The intraperitoneal and intravenous inoculation models, which involve the direct introduction of bacteria into the peritoneal cavity or bloodstream, respectively, allow for systemic dissemination of the bacteria and mimic severe scrub typhus and lethal outcomes(21, 23, 24). However, these routes bypass the natural mode of transmission, in which bacteria are introduced by the bite of an infected chigger mite and disseminate via the lymphatic system before bloodstream invasion(22, 25). The intradermal or subcutaneous mouse models, which better mimic the natural route of *Ot* infection, are commonly used to study bacterial dissemination and pathogenesis (15, 18, 19, 25–28). These models, though, are not effective for modeling severe scrub typhus disease. While intradermal infection leads to systemic bacterial dissemination and elicits host protective immune responses, immunocompetent mice (inbred C57BL/6 or outbred CD-1 mice, etc.) are largely resistant and develop only mild disease or no major symptoms(15, 19, 28). This suggests that mice are capable of mounting stronger protective immune responses against *Ot* than humans. Thus, developing a susceptible and immunocompetent mouse model is critical for investigating bacterial dissemination in humans, elucidating protective host immunity, and evaluating vaccine candidates against human relevant disease states.

It is known that *Ot* infection elicits strong type 1 immunity in both patients and animal models (29–32). We recently demonstrated that mice deficient in IFN-γ receptor signaling are highly susceptible to intradermal *Ot* infection, exhibiting 100% lethality even at very low infectious doses(25). In addition, these deficient mice display eschar-like lesions that are not observed in immunocompetent mice(25), highlighting the role of IFN-γ signaling in host protection against *Ot* infection. Although *Ifngr1*^-/-^ mice are highly susceptible to *Ot* infection, their immunocompromised status limits their ability to model the complex and coordinated immune responses observed during natural infections. As a result, they are not suitable for comprehensive studies of immunopathogenesis, vaccine development, or host protective immunity. Given that humans with intact IFN-γ signaling remain susceptible to *Ot*, we hypothesize that human susceptibility may stem from comparatively weaker IFN-γ signaling than that observed in mice. Although IFN-γ signaling is functionally conserved between humans and mice, species-specific differences may lead to differential regulation of downstream genes and contribute to the observed differences in susceptibility to infection (33, 34).

In this study, we establish a novel mouse model of scrub typhus by using a genetically engineered humanized mouse strain that lacks murine IFN-γ signaling but is reconstituted with functional human IFN-γ signaling. This immunocompetent mouse model exhibited increased susceptibility to *Ot* infection compared to wild-type (WT) B6 mice, as demonstrated by greater body weight loss, higher clinical disease scores, and elevated bacterial burdens in multiple organs. Further analysis demonstrated the impaired interferon-stimulated gene expression and dysregulated innate and adaptive immune cell responses in infected humanized mice. Notably, intradermal *Ot* infection in humanized mice resulted in eschar-like lesions that closely resemble those observed in human patients. Together, this humanized and immunocompetent mouse model, combined with intradermal infection, represents a promising platform for future scrub typhus studies on bacterial dissemination, immunopathogenesis, and vaccine development.

## Materials and Methods

### Ethics Statement

UTMB complies with the USDA Animal Welfare Act (Public Law 89–544), Health Research Extension Act of 1985 (Public Law 99–158), the Public Health Service Policy on Humane Care and Use of Laboratory Animals, and NAS Guide for the Care and Use of Laboratory Animals (ISBN-13). UTMB is registered as a Research Facility under the Animal Welfare Act and has current assurance with the Office of Laboratory Animal Welfare, in compliance with NIH policy. Infections were performed following Institutional Animal Care and Use Committee approved protocols (2101001A) at the UTMB in Galveston, TX.

### Animals, Infection, and Treatment

Male WT C57BL/6 (#000664) and *Ifngr1*^-/-^ (#003288) mice were obtained from the Jackson Laboratory. Humanized B-hIFNGR1/hIFNG/hIFNGR2 mice (#130949, C57BL/6 background) were purchased from Biocytogen (Waltham, MA). All mice were maintained under specific pathogen free conditions and challenged at 11-12 weeks of age. All mouse infection studies were performed in the ABSL3 facility in the Galveston National Laboratory located at UTMB; all tissue processing and analysis procedures were performed in the BSL3 or BSL2 facilities. All procedures were approved by the Institutional Biosafety Committee, in accordance with Guidelines for Biosafety in Microbiological and Biomedical Laboratories. For infection, mice were anesthetized using a VetFlo isoflurane vaporizer in an induction chamber. After anesthesia, the right flank was shaved using an electric trimmer to prepare the injection site. A 20 μL suspension of the *Ot* Karp strain (3×10^3^ FFU in SPG buffer) was injected intradermally into the right flank using a 0.3 mL insulin syringe with a 31G needle (Sol-Millennium, Chicago, IL). Mice were monitored for approximately 5 minutes (min) post-injection to ensure full recovery from anesthesia. Mice were monitored daily for body weight, skin eschar development and disease severity. At 14 days post-infection (dpi), mice were euthanized by CO inhalation, and tissues and blood were collected for further analysis.

### Bacterial Stock Preparation

Bacteria were inoculated onto L929 murine fibroblast monolayers in T150 cell culture flasks and gently rocked for two hours (h) at 37°C. After 2 h, Minimum Essential Medium with 10% fetal bovine serum, 100 units/mL of penicillin and 100 μg/mL of streptomycin were added. Cells were harvested by scraping at 7 dpi, re-suspended in Minimum Essential Medium, and lysed using 0.5 mm glass beads and vortexing for 1 min. The cell suspension was collected and centrifuged at 300×g for 10 min to pellet cell debris and glass beads. The supernatant from one T150 flask was further inoculated onto new monolayers of five T150 flasks. This process was repeated for a total of six passages. Cells from five flasks were pooled in a 50 mL conical tube with 20 mL medium and 5 mL glass beads. The conical tubes were gently vortexed at 10 sec intervals for 1 min to release the intracellular bacteria and placed on ice. The tubes were then centrifuged at 300×g for 10 min to pellet cell debris, and the supernatant was collected in Oakridge high speed centrifugation bottles, followed by centrifugation at 22,000×g for 45 min at 4°C to harvest bacteria. Sucrose-phosphate-glutamate buffer (0.218 M sucrose, 3.8 mM KH_2_PO_4_, 7.2 mM KH_2_PO_4_, 4.9 mM monosodium L-glutamic acid, pH 7.0) was used for preparing bacterial stocks, which were stored at −80°C(24, 35, 36). The same lot of stocks were used for all experiments described in this study. The titers of the bacterial stocks were measured by focus forming assays, as previously described(21).

### qPCR for Measuring Bacterial Burdens

Bacterial burdens were calculated from collected mouse tissues and cultured cells. The samples were initially incubated with proteinase K and lysis buffer at 56°C overnight. DNA was extracted using the DNeasy Blood and Tissue Kit (Qiagen, Germantown, MD) according to the instructions and used for qPCR assays as previously described(21, 26, 37, 38). The 47-kDa gene was amplified using the primer pair OtsuF630 (5’-AACTGATTTTATTCAAACTAATGCTGCT-3’) and OtsuR747 (5’-TATGCCTGAGTAAGATACGTGAATGGAATT-3’) primers (IDT, Coralville, IA) and detected with the probe OtsuPr665 (5’-6FAM-TGGGTAGCTTTGGTGGACCGATGTTTAATCT-TAMRA) (IDT, Coralville, IA) by SsoAdvanced Universal Probes Supermix (Bio-Rad, Hercules, CA). Bacterial burdens were normalized to total microgram (μg) of DNA per μL for the same samples. Absolute quantification was performed using a 10-fold serial dilution of the *Ot* Karp 47-kDa plasmid carrying a single copy of the target gene.

### Quantitative Reverse Transcriptase PCR (qRT-PCR)

RNA was extracted from mouse lungs and brain using the RNeasy Mini Kit (Qiagen, Germantown, MD) according to the manufacturer’s instruction, followed by the quality and quantity assessment using a BioTek microplate reader. cDNA was synthesized by using the iScript Reverse Transcription kit (Bio-Rad, Hercules, CA) with the same amount of RNA (1 μg). qRT-PCR was performed using 5 μL of iTaq SYBR Green Supermix (Bio-Rad, Hercules, CA), 1 μL of a forward and reverse primer mix (0.5 μM final concentration of each), and 4 μL of diluted cDNA. Samples were denatured for 30 s at 95°C, followed by 40 cycles of 15 s at 95°C, and 60 s at 60°C, utilizing a CFX96 Touch real-time PCR detection system (Bio-Rad, Hercules, CA).

Relative quantitation of transcript levels was calculated using the 2−ΔΔCt method and normalized to glyceraldehyde-4-phosphate dehydrogenase (*Gapdh*). All primers were generated by Integrated DNA Technologies (IDT, Coralville, IA) and are listed in **S1 Table**. The primer sequences were obtained from PrimerBank (https://pga.mgh.harvard.edu/primerbank).

### Bio-Plex Assay

Whole blood was collected from euthanized mice at 14 dpi, and serum was isolated by using serum separator tubes (BD Bioscience, San Diego, CA). Bacterial inactivation was performed on the samples, as described in our previous study(21). The Bio-Plex Pro Mouse Cytokine Th1 Panel was used to measure serum cytokine levels. The Bio-Rad Bio-Plex Plate Washer and Bio-Plex 200 machine located in the UTMB Flow Cytometry and Cell Sorting Core Lab were used for sample processing and analysis.

### Histology

Tissues were fixed in 10% neutral buffered formalin and embedded in paraffin at the UTMB Research Histology Service Core. Tissue sections (5-μm thickness) were stained with hematoxylin and eosin and mounted on slides. Sections were imaged under an Olympus BX53 microscope, and at least five random fields for each section were captured.

### Clinical Pathology

Animal blood chemistry analysis was performed by using the VetScan Chemistry Analyzer (Zoetis, Parsippany-Troy Hills, NJ), according to the manufacturer’s instruction. Briefly, mouse serum (100 μL) was uploaded into the VetScan Comprehensive Diagnostic Profile reagent rotor, which was used for quantification of alanine aminotransferase (ALT), albumin (ALB), alkaline phosphatase (ALP), amylase (AMY) total calcium (CA^++^), globulin (GLOB), glucose (GLU), phosphorus (PHOS), potassium (K^+^), sodium (NA^+^), total protein (TP), and urea nitrogen (BUN). This analysis was performed at the UTMB ABSL3 animal facility.

### ELISA

Mouse serum samples were collected and human IFN-γ concentrations were analyzed using the Legend Max Human IFN-γ ELISA Kit (Biolegend, San Diego, California) according to the manufacturer’s instructions provided by the vendor. To test total IgM and IgG titers in the serum, 96-well plates were precoated with 2 μg/mL of recombinant *Ot* Karp strain TSA56 protein in PBS and blocked with 0.5% BSA. The recombinant TSA56 protein was generated by Genscript (Piscataway, NJ). Serum was diluted 1:3 until endpoint titers were determined. Goat anti-mouse IgM-HRP (1021-05, Southern Biotech) and goat anti-mouse IgG-HRP (1030-05, Southern Biotech) were used at a 1:3000 dilution for detection. 1-Step Ultra TMB ELISA Substrate Solution (Thermo Fisher Scientific) was used as the visualizing reagent. Optical density was measured using a Bio-Tek microplate reader. Area under the curve analysis was performed on each curve at every timepoint.

### Flow Cytometry

Spleen single-cell suspensions were prepared by passing spleen tissues through 70-μm cell strainers. Red blood cells were removed by using Red Cell Lysis Buffer (Sigma-Aldrich, St. Louis, MO) for 5 min at RT. For surface marker analysis, leukocytes were stained with the Fixable Viability Dye (eFluor 506, Thermo Fisher Scientific, Waltham, MA) for live/dead cell discrimination, blocked with FcγR blocker, and incubated with fluorochrome-labeled antibodies for 30 min. The fluorochrome-labeled antibodies were purchased from Thermo Fisher Scientific and Biolegend as below: Alexa Fluor 700 anti-CD11b (M1/70), APC anti-Ly6G (1A8), PE/Dazzle-594 anti-Ly6C (HK1.4), Percp-Cy5.5 anti-CD11c (N418), FITC anti-MHCII (M5/114.15.2), PE-Cy7 anti-CD3ε (145-2C11), Percp-cy5.5 anti-CD4 (GK1.5), APC-Cy7-anti-CD8a (53–6.7), BV711 anti-CD44 (IM7), APC anti-CD62L (MEL-14), PE CF594-anti-NK1.1 (PK136), FITC anti-CD69 (H1.2F3) and Percp-Cy5.5 CTLA4 (UC10-4B9). BV421 anti-F4/80 (T45-2342) antibody was purchased from BD Bioscience (San Diago, CA). For Foxp3 staining, cells were fixed and permeabilized using Foxp3 / Transcription Factor Fixation/Permeabilization kit (Thermo Fisher Scientific), followed by the staining with PE anti-Foxp3 (FJK-16s) for 45 min. Cell samples were fixed in 2% paraformaldehyde overnight at 4°C, acquired by a BD Symphony A5 and analyzed via FlowJo software version 10 (BD, Franklin Lakes, NJ). The gating strategy of immune cell subsets is based on recent publications(25, 37, 39).

### Statistical Analysis

Data are presented as mean ± standard deviation (SD). The data related to qRT-PCR, disease scores, skin lesion scores, flow cytometry, Bio-Plex and chemistry parameters were analyzed with one-way ANOVA. Bacterial burdens between two groups were analyzed by the student t-test. Body weight changes were analyzed by two-way ANOVA. After a significant F-test for the ANOVA model, either Tukey’s multiple comparisons test or Šídák’s multiple comparisons test was used for comparison between groups. All data were analyzed by using GraphPad Prism software 10. Statistically significant values are denoted as * *p* < 0.05, ** *p* < 0.01, *** *p* < 0.001, and **** *p* < 0.0001, respectively.

## Results

### Humanized mice are susceptible to *Ot* infection

Our previous study demonstrated that IFN-γ signaling is essential for host defense against *Ot* infection (25). Given that human IFN-γ signaling may be less robust than its murine counterpart(34), we hypothesized that genetically engineered mice carrying humanized IFN-γ signaling may exhibit increased susceptibility to *Ot* infection. To test this, we utilized a genetically engineered humanized mouse model that possesses functional human IFN-γ signaling while lacking murine IFN-γ signaling. WT and *Ifngr1*^-/-^ mice were included as resistant and highly susceptible immunological control models, respectively(25). We chose to use the intradermal infection route, as established in our previous studies(25, 26), to closely mimic the natural mode of *Ot* transmission. As shown in **Figure 1A and B**, WT mice showed marginal body weight changes during infection with low disease scores. However, *Ifngr1*^-/-^ mice were of high susceptibility, exhibiting a consistently decreasing body weight and increasing disease scores at 11 and 14 dpi. Humanized mice also showed similar body weight changes and disease scores as *Ifngr1*^-/-^ mice until 11 dpi, but these mice started to gain body weight at 12 dpi and continued to recover. By 14 dpi, humanized mice exhibited a comparable disease score as WT mice. Previously, we have shown the first report that *Ifngr1*^-/-^ mice, but not WT mice, developed an eschar lesion at the skin inoculation site, which was characterized by necrosis and the formation of a black crust with a red halo, as seen in scrub typhus patients(25). In this study, we observed that humanized mice also formed skin eschar lesions of comparable sizes as *Ifngr1*^-/-^ mice at 11 dpi (**Figure 1C and D**). H&E staining of skin tissues in *Ifngr1*^-/-^ and humanized mice revealed extensive inflammatory cell infiltration. No eschar lesion and immune cell infiltration were found in skin samples of uninfected mice at 14 dpi (**S2 Fig**); WT mice exhibited minimal immune cell infiltration (**Figure 1E**). Therefore, our study provides the first evidence of skin eschar formation resembling human scrub typhus, and heightened susceptibility to *Ot* infection using an immunocompetent humanized mouse model.

**Figure 1.**
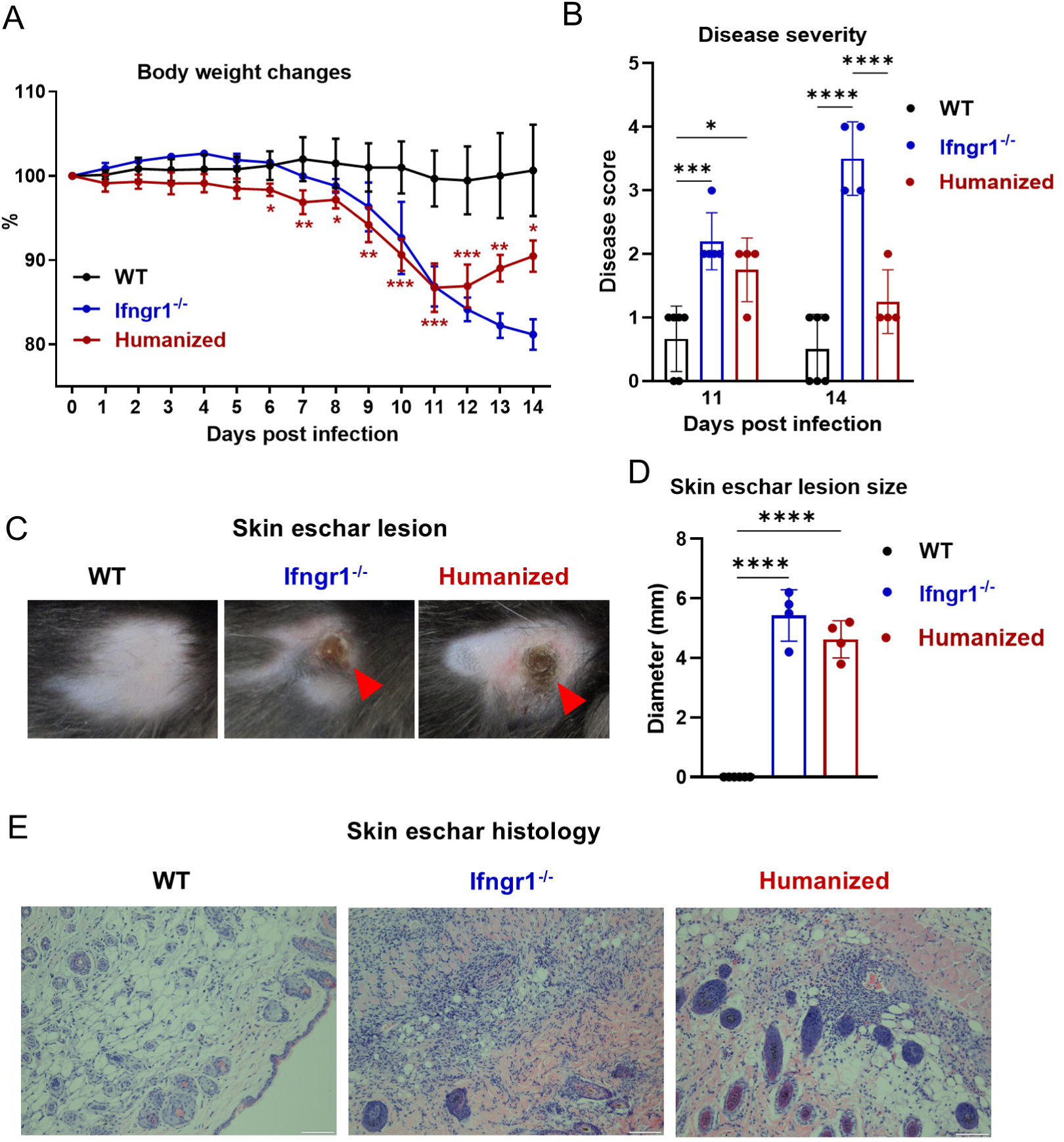
Humanized mice are susceptible to *Ot* infection. WT B6 (n=6), *Ifngr1*^-/-^ (n=4) and humanized (n=4) mice were intradermally infected with *Ot* (3×10^3^ FFU) on the right flank. Mice were monitored daily for (A) body weight changes and (B) disease scores. (C) Skin eschar lesions at inoculation sites at 11 dpi are presented and (D) their sizes are measured. (E) Representative histological images of the skin eschar lesion are shown at 11 dpi. Scale bar = 100 µm. Data are presented as mean ± SD from two independent pooled experiments. Body weight changes were analyzed by two-way ANOVA with Tukey’s multiple comparisons between WT and humanized mice at indicated time points. Disease scores (11 and 14 dpi) and skin eschar lesion size (11 dpi) were analyzed by one-way ANOVA with Tukey’s multiple comparisons. *, *p* < 0.05; **, *p* < 0.01; ***, *p* < 0.001; ****, *p* < 0.0001.

### Systemic inflammation in humanized mice following *Ot* infection

To assess whether *Ot* infection induces inflammatory responses in major organs of humanized mice, we performed histological analyses on the liver, lung, and brain tissues at 14 dpi (**Figure 2**). In uninfected WT and humanized mice, no notable immune cell infiltration was detected in the liver. Following *Ot* infection, WT mice showed mild inflammatory cell infiltration limited to the portal triads, with minimal involvement of the lobular regions. In contrast, humanized mice exhibited moderate portal inflammation accompanied by inflammatory cell infiltration extending into the lobular areas. No tissue necrosis was observed in any samples. For the lung tissues, uninfected samples saw intact alveolar architecture, accompanied by minimal or no cellular infiltration. The lung histology of infected humanized mice revealed increased mononuclear cell infiltration throughout the interstitium compared to infected WT mice, suggesting moderate interstitial pneumonitis in infected humanized mice. It is known that *Ot* infection can cause blood brain barrier disruption and central nervous system disorders(35, 40, 41). We also found that the meningeal layer of humanized mice with *Ot* infection showed clear mononuclear cell infiltration, indicating meningitis or meningoencephalitis. Overall, our data demonstrated systemic inflammation across multiple organs in the humanized mouse model of scrub typhus.

**Figure 2.**
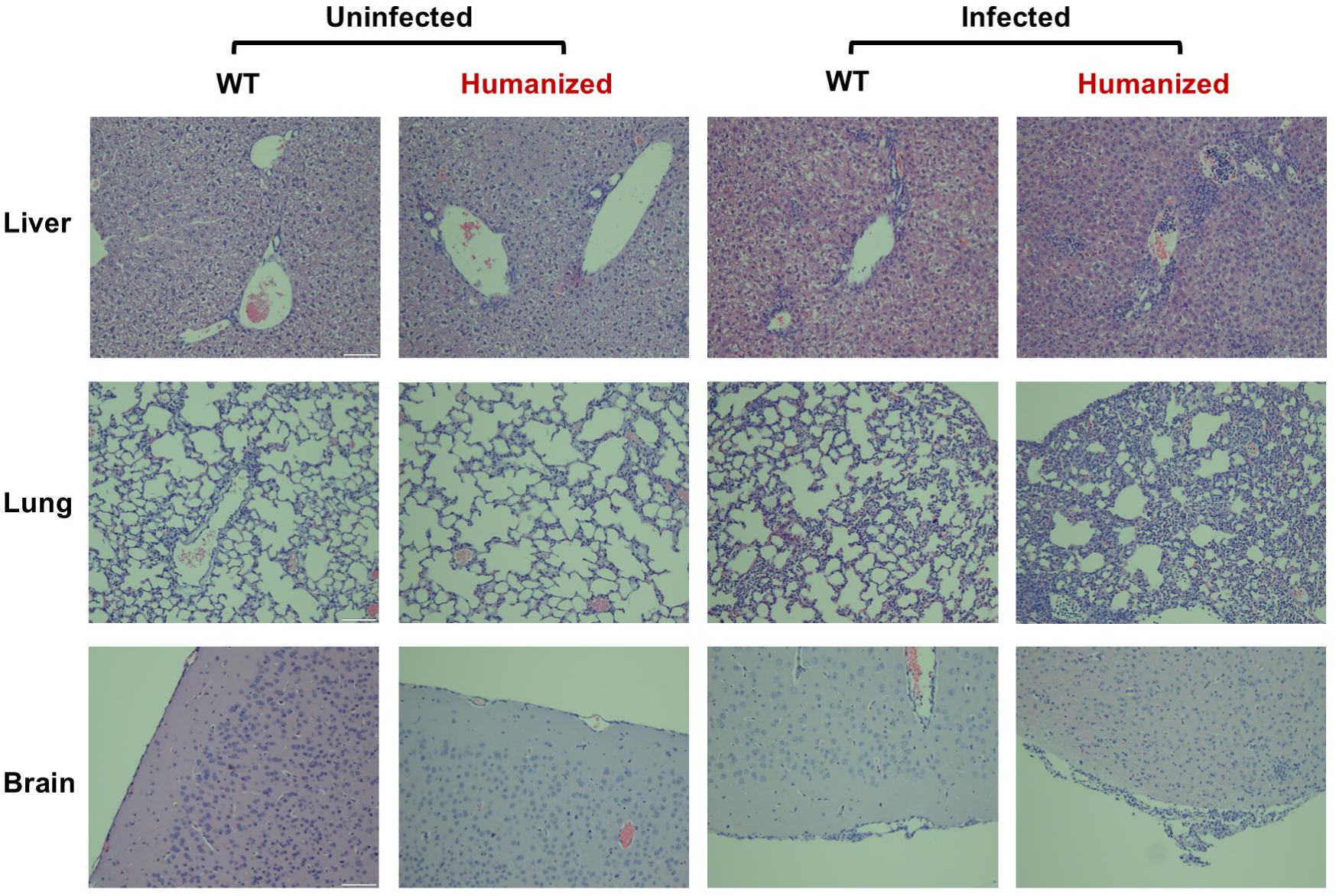
Tissue histology of WT, *Ifngr1*^-/-^ and humanized mice following *Ot* infection. WT B6 (n=9) and humanized (n=6) mice were infected as described in Figure 1. Uninfected WT B6 (n=3) and humanized (n=2) were i.d. injected with L929 cell culture control. Representative histological images of the liver, lung and brain are shown at 14 dpi. Scale bar = 100 µm.

### *Ot* infection induces human, but not murine IFN-**γ** signaling in humanized mice

To confirm the activation of human IFN-γ signaling in humanized mice following infection, we measured the transcript levels of *Ifngr1* and *Ifngr2* in brain and lung tissues, which are the primary *Ot* target organs for severe disease outcomes(42). As shown in **Figure 3A** and **B**, WT mice expressed mouse, but not human *Ifngr1* and *Ifngr2* in both the brain and lungs, while humanized mice only expressed human but not mouse *Ifngr1* and *Ifngr2*. We further quantified human IFN-γ protein levels in sera by using specific ELISA. We found that human IFN-γ levels in the serum were minimal in uninfected humanized mice but increased significantly after infection (**Figure 3C**), and that human IFN-γ was undetectable in uninfected or infected WT mice. Altogether, our results demonstrated that humanized mice activate human but not mouse IFN-γ responses after *Ot* infection.

**Figure 3.**
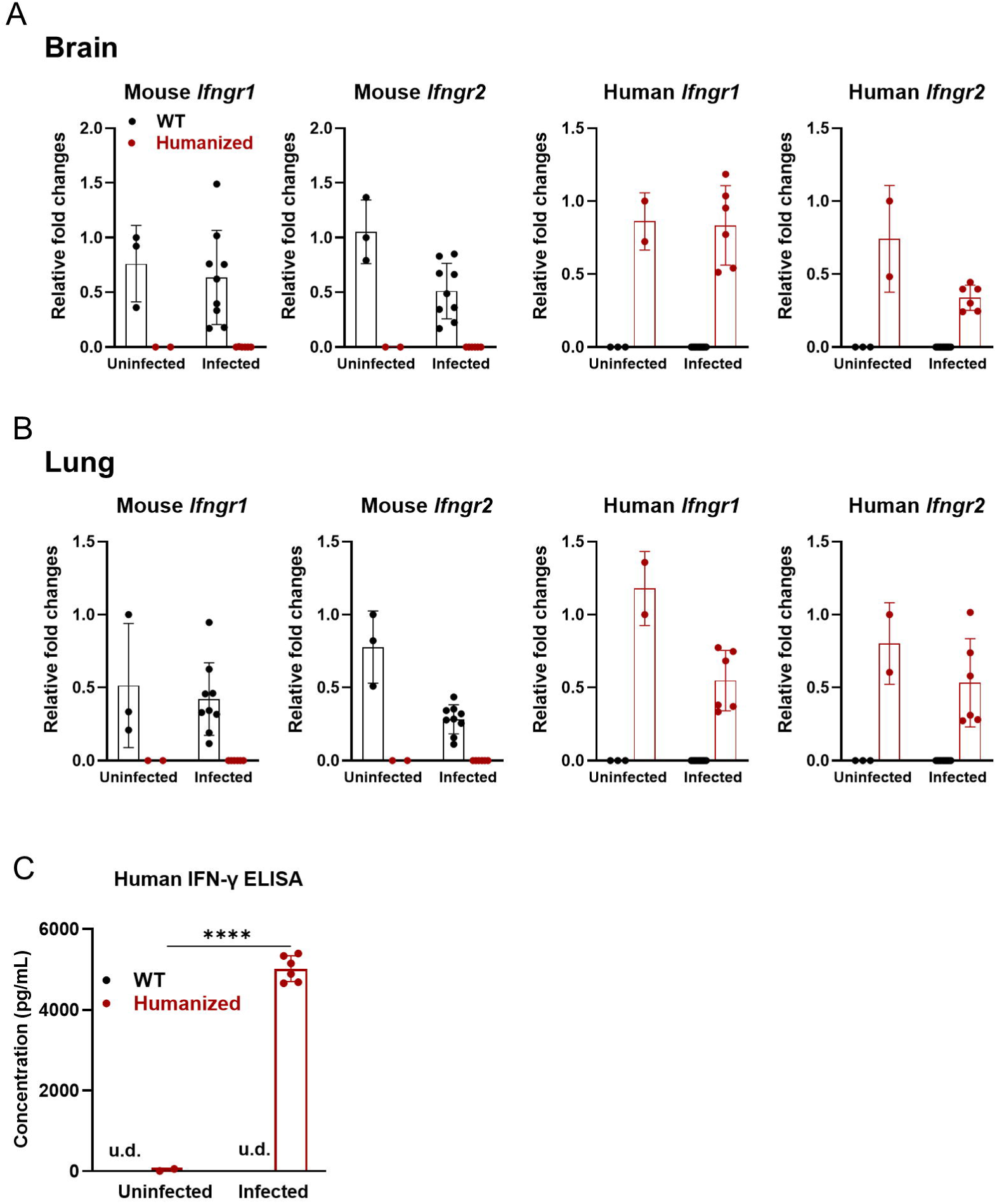
Expression of *Ifng*, *Ifngr1* and *Ifngr2* in WT and humanized mice. Mice were infected as described in Figure 2. (A-B) RNA was extracted from mouse brain and lung tissues. Mouse and human *Ifngr1* and *Ifngr2* transcript levels were analyzed by RT-qPCR, and the relative fold changes are shown. (C) Human IFN-γ levels in the sera of WT B6 and humanized mice were measured by sandwich ELISA. The label “u.d.” represents undetectable levels. Data are presented as mean ± SD from three independent pooled experiments.

### Serum cytokines and biochemical profiles in humanized mice with *Ot* infection

To better understand the status of systemic immune responses, we analyzed serum cytokines of WT and humanized mice by Bio-Plex assay. Reinforcing the RT-qPCR data of **Figure 3**, the Bio-Plex assay showed that mouse IFN-γ was significantly upregulated in infected WT mice, but it was undetected in humanized mice (**Figure 4**). A general trend of cytokine upregulation was observed in both WT and humanized mice following infection, with increased levels of IL1β, IL-2, IL-10 and IL-6. Of these, serum IL-6 was significantly elevated in infected-humanized mice when compared to WT controls, suggesting a heightened IL-6-mediated inflammatory response in the humanized mouse model.

**Figure 4.**
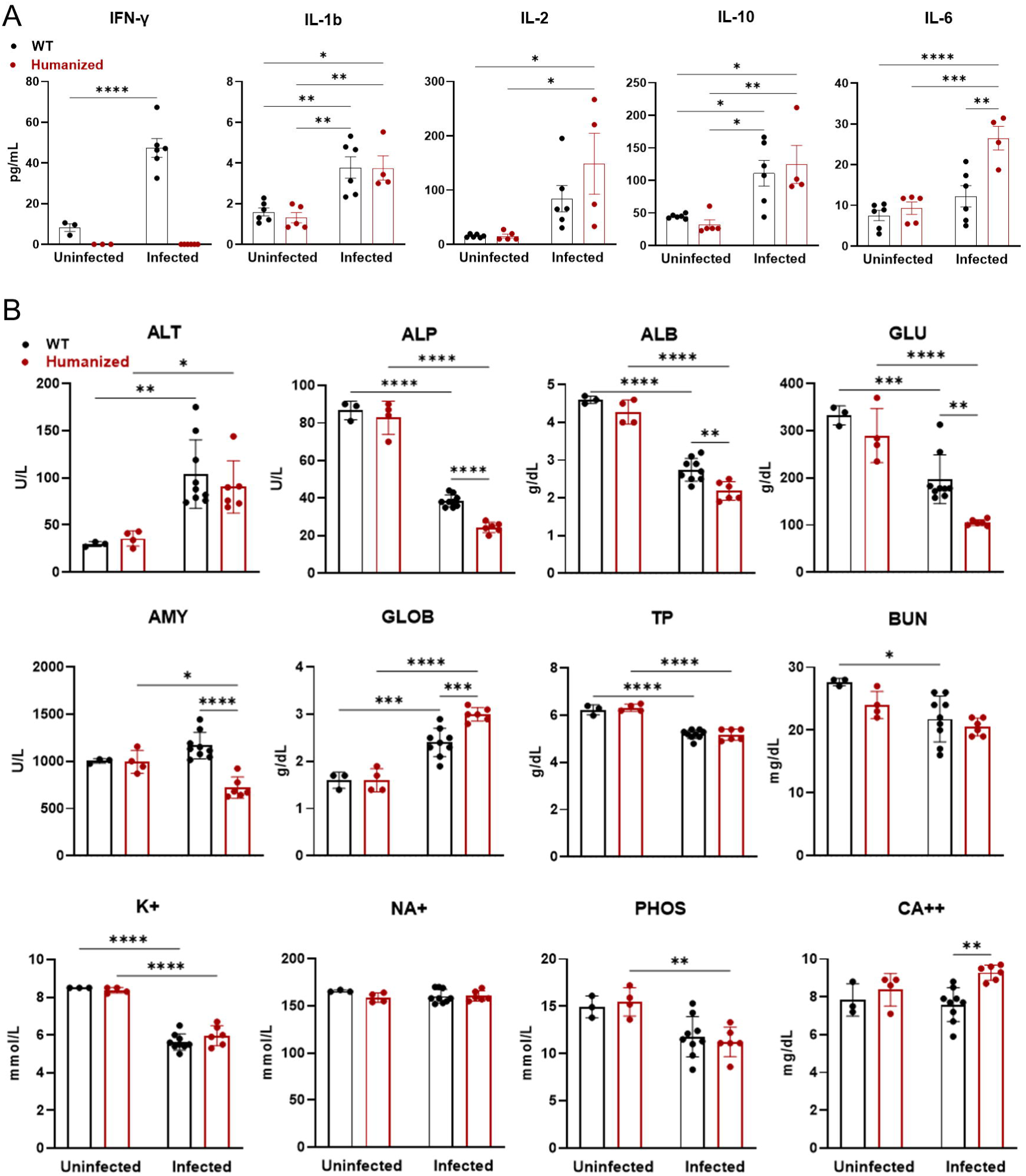
Serum cytokines and chemistry profiles in humanized mice with *Ot* infection. Mice were infected as described in Figure 1. (A) Serum cytokine and chemokine levels were analyzed at 14 dpi by a Bio-Plex assay. (B) The serum chemistry parameters were detected by using VetScan Comprehensive Diagnostic Profile Reagent Rotor. The parameters include alanine aminotransferase (ALT), albumin (ALB), alkaline phosphatase (ALP), amylase (AMY), total calcium (CA^++^), globulin (GLOB), glucose (GLU), potassium (K^+^), sodium (NA^+^), total protein (TP), blood urea nitrogen (BUN), and phosphorus (PHOS). Data are shown as mean ± SD from three pooled independent experiments and analyzed by one-way ANOVA with a Tukey’s multiple comparisons test. *, *p* < 0.05; **, *p* < 0.01; ***, *p* < 0.001; ****, *p* < 0.0001.

To evaluate animal physiological status, we measured serum biochemical parameters by using VetScan Comprehensive Diagnostic Profile reagent rotors. In the absence of infection, humanized mice showed no physiological abnormalities, as indicated by serum biochemical parameters comparable to those of WT mice. (**Figure 4B**). However, *Ot* infection resulted in significantly decreased levels of alkaline phosphatase (ALP), albumin (ALB), and glucose (GLU), with more pronounced reductions observed in humanized mice. Amylase (AMY) levels showed a significantly increased trend in WT mice after infection, whereas a more pronounced decrease was observed in humanized mice, consistent with findings in *Ifngr1* / mice(25). These results may suggest compromised liver function and malnutrition in humanized mice. This pattern of biochemical changes closely aligns with our previous observations in *Ifngr1*-deficient mice(25). In addition, a further increase of globulin (GLOB) in humanized mice may indicate heightened inflammation in response to infection. Both WT and humanized infected mice demonstrated a dysregulation of total protein (TP), alanine aminotransferase (ALT), phosphorus (PHOS) and potassium (K+). These findings suggest an enhanced systemic inflammatory response and disrupted physiological homeostasis in humanized mice following *Ot* infection.

To assess whether humanized mice exhibit altered humoral immunity in response to *Ot* infection, we measured *Ot*-specific antibody titers in mouse sera. ELISA results showed comparable levels of IgM and IgG against *Ot* TSA56 antigen between humanized and WT mice (**S3 Fig**), indicating that humanized mice mount a competent humoral immune response like WT controls.

### Elevated bacterial burden but reduced interferon-stimulated gene (ISG) expression in humanized mice

To assess bacterial control in humanized mice, bacterial burdens were measured in various organs (**Figure 5A**). Consistent with previous findings(22, 25), the highest bacterial burden was observed in mouse lungs, followed by the spleens in both WT and humanized mice. We found that humanized mice exhibited approximately a 2-fold increase in bacterial burden in the lungs and spleen. Notably, a 10-fold increase of bacterial burdens in the brain was detected in humanized mice as compared to WT mice. Likewise, the kidney and liver organs in humanized mice also showed significantly higher bacterial burden. These results suggest that human IFN-γ signaling confers weaker antibacterial activity compared to its murine counterpart.

**Figure 5.**
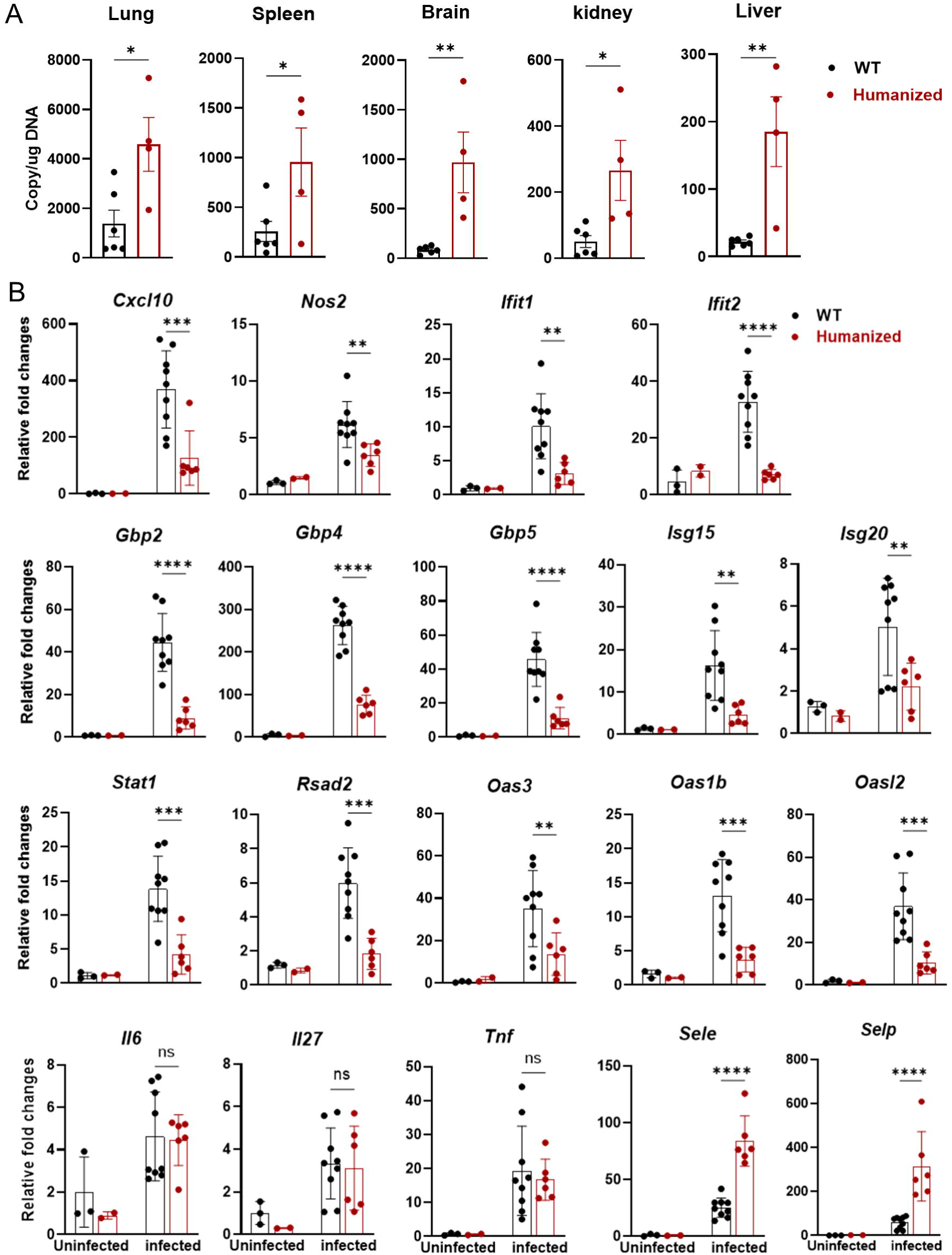
Humanized mice exhibited an increased bacterial burden and reduced expression of interferon stimulated genes (ISGs) following *Ot* infection. Mice were infected as described in Figure 1 and were euthanized at 14 dpi. (A) Genomic DNA was isolated from the lung, spleen, brain, kidney and liver. Bacterial burden was measured by qPCR. (B) The transcript levels of ISGs in the brain were analyzed by qRT-PCR. Data are shown as mean ± SD from three pooled independent experiments and analyzed by unpaired t-test for bacterial burdens and one-way ANOVA with a Šídák’s multiple comparisons test for ISG expression. *, *p* < 0.05; **, *p* < 0.01; ***, *p* < 0.001; ****, *p* < 0.0001; ns, no significant difference.

IFN-γ signaling activates the JAK-STAT pathway and induces the expression of multiple downstream ISGs, driving a robust antibacterial response(43). Disruption of IFN-γ/STAT1 signaling has been shown to result in uncontrolled bacterial replication and lethal *Ot* infection(25). To assess the antibacterial response of the humanized scrub typhus model, we examined ISG transcript levels in various organs by qRT-PCR. Our results demonstrated that as compared to WT mice, humanized mice showed lower expression of all examined ISGs in the brain, including *Cxcl10*, *Nos2*, *Ifit1/2*, *Gbp2/4/5*, *Isg15*, *Isg20*, *Stat1*, *Rsad2*, *Oas3*, *Oas1b* and *Oasl2* (**Figure 5B**). Furthermore, we confirmed that the lungs of humanized mice also exhibited lower expression of some ISGs including *Cxcl10*, *STAT1*, *Ifit1/2*, *Rsad2*, *Isg15,* and *Oasl2* (**S4 Fig**). In addition, the inflammatory cytokine levels (*IL6*, *Tnf* and *Il27)* were comparable in both lungs and brain between two groups (**Figure 5B and S4 Fig**). Interestingly, we found that humanized mice expressed significantly higher levels of *Sele* (E-selectin) and *Selp* (P-selectin) in the brain, but not in the lungs, indicating increased endothelial activation and inflammation in the brain of humanized mice with *Ot* infection.

### Dysregulated innate and adaptive immune cell response in humanized mice following *Ot* infection

*Ot* infection triggers robust innate and adaptive immune responses, characterized by strong type 1 immunity and activation of multiple immune cell subsets, including NK cells, macrophages, dendritic cells, neutrophils, and effector T cells. While these cells play critical roles in controlling bacterial replication and dissemination, their excessive activation can lead to immunopathogenesis. We found that *Ot* infection induced stronger CD4^+^ and CD8^+^ T effector cell responses in the spleens of humanized mice compared to WT controls (**Figure 6A**). However, there was a significantly decreased percentage of Treg cells among CD4^+^ T cells (**Figure 6B**). Moreover, CTLA4 Treg cell numbers were markedly reduced in humanized mouse spleens (**Figure 6C**). This result suggests that *Ot* infection leads to excessive T cell activation accompanied by a diminished immunosuppressive T cell response in humanized mice.

**Figure 6.**
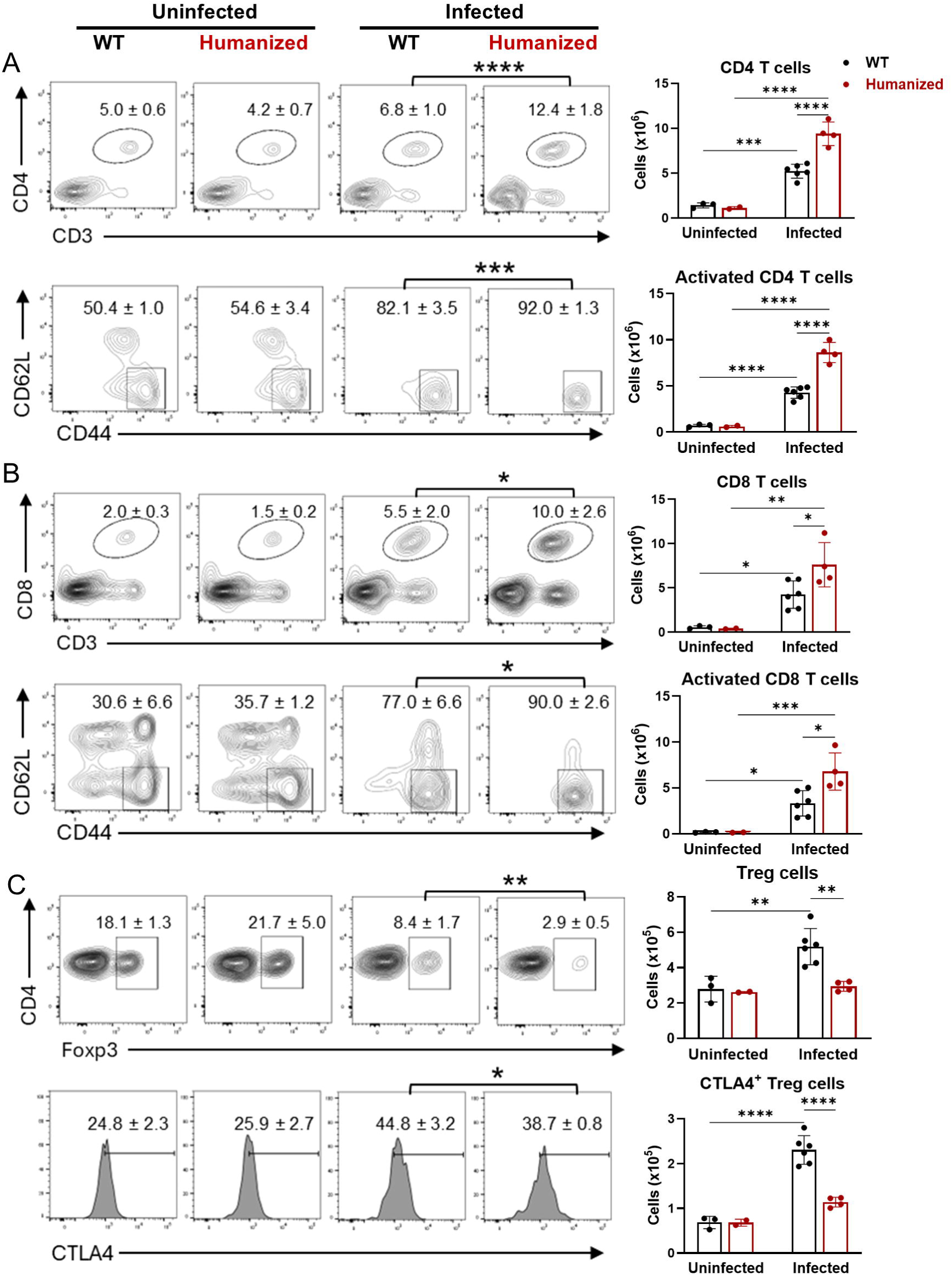
Imbalanced T cell responses in humanized mice following infection. Mice were infected as described in Figure 1 and were euthanized at 14 dpi. Spleen cells were isolated and analyzed by flow cytometry. Single cells were gated first by FSC/SSC, followed by the exclusion of dead cells by live/dead staining. (A-B) CD4^+^ and CD8^+^ T cells were gated on CD3^+^CD4^+^ and CD3^+^CD8^+^, respectively. CD44^+^CD62L^-^ T cells were identified as activated populations. (C) For regulatory T (Treg) cells, Foxp3 was gated on CD4^+^ T cells and the expression of CTLA4 was further gated on CD4^+^Foxp3^+^ T cells. The percentages of cell populations were shown as mean ± SD on the flow cytometric images and the statistical analysis between infected WT and humanized mice were labeled. The absolute numbers of cell populations were also calculated and shown next to the flow cytometric images. One-way ANOVA with a Šídák’s multiple comparisons test was used for data analysis. *, *p* < 0.05; **, *p* < 0.01; ***, *p* < 0.001; ****, *p* < 0.0001.

### Excessive neutrophil response and limited innate immune cell activation in humanized mice

Neutrophilia is observed in scrub typhus patients and is linked to disease progress(44, 45). We found that infected humanized mice exhibited 3.4-fold higher neutrophil numbers in the spleen as compared to WT controls (**Figure 7A**). Monocytes were also recruited into the spleen following infection, with comparable infiltration levels observed between WT and humanized mice (**Figure 7B**). Notably, both macrophages and dendritic cells in infected humanized mice exhibited reduced MHC II expression, indicating impaired activation of antigen-presenting cells in response to human IFN-γ signaling. Humanized mice also harbored significantly fewer NK cells, which displayed a markedly lower activation status, as evidenced by reduced CD69 expression (**Figure 7C**). Overall, our results suggest an increased neutrophil response alongside impaired activation of other innate immune cells in humanized mice of *Ot* infection.

**Figure 7.**
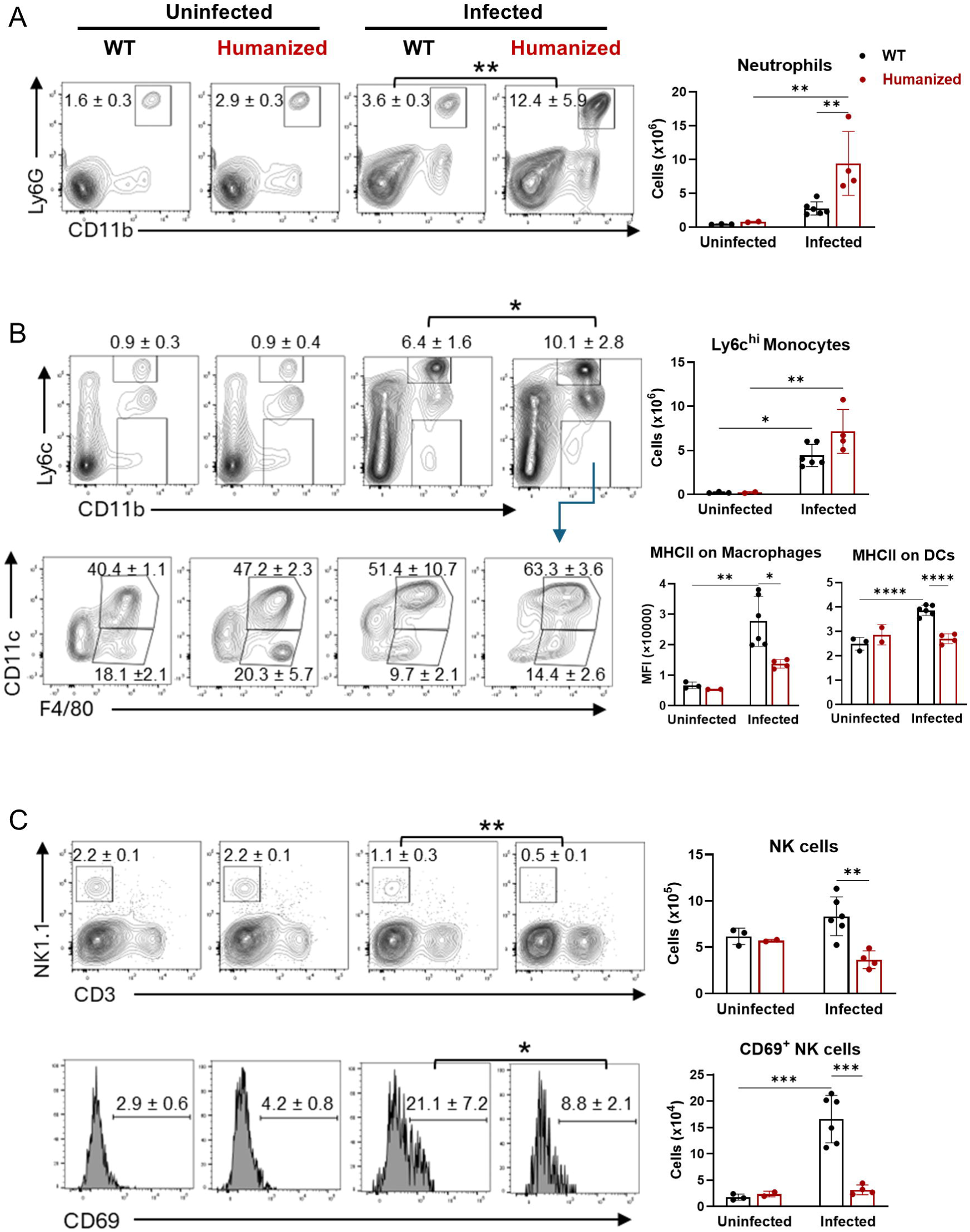
Excessive neutrophils and reduced innate immune cell activation in humanized mice following *Ot* infection. Mice were infected as described in Figure 1 and were euthanized at 14 dpi. Spleen cells were isolated and analyzed by flow cytometry. Single cells were gated first by FSC/SSC, followed by the exclusion of dead cells by live/dead staining. (A-B) Neutrophils and monocytes were identified by CD11^+^Ly6G^+^ and CD11b^+^Ly6C^hi^, respectively. CD11b^+^Ly6C^-^ cells were further gated by CD11c and F4/80 markers. Macrophages and dendritic cells were characterized by CD11c^-^F4/80^+^ and CD11c^hi^F4/80^+^, respectively. The mean fluorescent intensity (MFI) of MHCII on macrophages and dendritic cells were measured. (C) NK cells were gated on CD3^-^NK1.1^+^, and CD69 was used as an activation marker of NK cells. The percentages of cell populations were shown as mean ± SD on the flow cytometric images and the statistical analysis between infected WT and humanized mice were labeled. The absolute numbers and MFI of cell populations were also calculated and were shown next to the flow cytometric images. One-way ANOVA with a Šídák’s multiple comparisons was used for data analysis. *, *p* < 0.05; **, *p* < 0.01; ***, *p* < 0.001; ****, *p* < 0.0001.

## Discussion

Scrub typhus is a systemic, life-threatening disease with a high incidence in the “tsutsugamushi triangle”, yet it remains significantly neglected(46). A better understanding of *Ot* pathogenesis and the development of effective vaccines represent the most promising strategies for preventing future outbreaks of this disease(47, 48). Although several animal models have been established by other researchers and our group(14–19, 21, 22, 28, 49, 50), there remains a need for an accurate mouse model that closely recapitulates human scrub typhus. In this study, we introduce a newly developed genetically engineered humanized mouse model for scrub typhus, featuring functional human IFN-γ signaling in the absence of murine IFN-γ signaling. We demonstrated that this humanized mouse strain exhibits increased susceptibility to intradermal *Ot* infection, characterized by multiorgan bacterial dissemination, skin eschar formation, systemic inflammation, altered serum biochemical profiles, and dysregulated innate and adaptive immune responses. This study represents the first establishment of a scrub typhus model using genetically engineered humanized mice, and our findings support the utility of this model as a promising tool for future studies on pathogenesis, drug screening, and vaccine development.

We have previously demonstrated that IFN-γ signaling is necessary for host defense against *Ot* infection as the *Ifngr1*^-/-^ mouse strain, in contrast to WT B6 mice, develops a lethal model of scrub typhus and forms a skin eschar lesion at the site of intradermal inoculation(25). The eschar serves as a key diagnostic indicator in patients with acute febrile illness in regions endemic for scrub typhus(51–53). Clinical observation indicates that the presence of an eschar might be associated with increased disease severity(54, 55). Our study reports skin eschar lesions in an immunocompetent mouse model in response to *Ot* infection for the first time. Representative HE staining of eschar lesions show apparent immune infiltration in both humanized and *Ifngr1*^-/-^ mice, but not in WT mice (Figure 1E). Therefore, our data generated from our humanized mouse model (Figure 1), along with the previously published *Ifngr1*^-/-^ model(25) suggest that the species-specific differences in IFN-γ signaling might be a critical determinant of eschar formation and disease severity of scrub typhus. The downstream mechanisms by which IFN-γ signaling influences eschar formation, however, still remain unclear. Interestingly, unlike mouse cells that express strong inducible nitric oxide synthase (iNOS) by IFN-γ stimulation, human cells instead produce indoleamine dioxygenase (34). Due to the key role of iNOS in wound healing(56), it is possible that compromised IFN-γ/iNOS signaling contributes to eschar formation in not only scrub typhus, but also rickettsial infection(57). Further studies are needed to elucidate downstream mechanisms by which IFN-γ signaling influences eschar formation.

Scrub typhus can cause systemic infection and multiorgan dysfunction, particularly in cases with delayed diagnosis and treatment. Immune infiltration in response to infection leads to inflammatory damage and organ failure in patients(52, 58–60). Our histology assessment, serum Bio-plex assay, biochemical analysis and bacterial burden measurement collectively demonstrate systemic bacterial dissemination and inflammatory responses across multiple organs, including lung, liver and brain in the humanized mouse model (Figure 2, 4 and 5). The immune and biochemical profiles in humanized mice revealed unique features consistent with those observed in our recently published *Ifngr1*^-/-^ mouse model(25). For example, serum analysis revealed elevated IL-6 in humanized mice compared to WT mice following infection, a characteristic that is also noted in scrub typhus patients (61, 62). Similar IL-6 upregulation was also observed in *Ifngr1*^-/-^ mice infected with *Ot*(25), indicating that the proinflammatory cytokine IL-6 may serve as a potential biomarker for predicting disease severity in scrub typhus. In addition, the decreased serum levels of ALP, ALB and GLU which were found in infected *Ifngr1*^-/-^ mice(25), were also observed in humanized mice compared to WT controls (Figure 4), indicating excessive liver dysfunction in this novel mouse model. Therefore, combining serum cytokine profiles and biochemical parameters with bacterial load may offer a more accurate prediction of disease severity in scrub typhus patients.

Scrub typhus has become a leading cause of central nervous system infection in endemic regions(63, 64). Severe infection can lead to a range of neurological complications, including meningitis, encephalitis, and meningoencephalitis, which are associated with increased mortality(35, 41, 65–68). Moreover, *Ot* infection may increase the risk of long-term sequelae and neurodegenerative diseases, such as dementia(69, 70). In this study, the increased susceptibility of humanized mice to *Ot* infection is evidenced by higher bacterial burdens in multiple organs compared to WT mice (Figure 5A). Notably, brain tissues of humanized mice exhibited the most pronounced increase, with bacterial burdens approximately 10-fold higher than those in WT controls. The uncontrolled bacterial replication observed in the brain may be attributed to the substantial downregulation of several ISGs (Figure 5B). Consistent with this, histological analysis also showed significant immune cell infiltration within the brain meninges of humanized mice, which may contribute to meningoencephalitis (Figure 2). The significantly elevated expression of key adhesion molecules *Sele* (E-selectin) and *Selp* (P-selectin) in the brains of humanized mice may indicate robust neuroinflammation and potential blood-brain barrier dysfunction(71–73). These findings may support the utility of the humanized mouse model as a valuable tool for investigating neurological complications associated with scrub typhus.

We further evaluated innate and adaptive immune responses in humanized mice following infection (Figure 5-7, S2 Fig and S3 Fig). Activation of IFN signaling induces robust expression of ISGs, which play a critical role in controlling pathogen replication(74–76). Thus, we first analyzed a comprehensive ISG expression panel in both the brain and lungs. We found that the expression of several ISGs (*Ifit1*, *Ifit2, Rsad2, Isg15, Cxcl10* and *Oasl2*) were downregulated in both organs of humanized mice, whereas members of the *Gbp* family (*Gbp2*, *Gbp4* and *Gbp5*), were selectively reduced in the brain. Given that bacterial burdens in humanized mice were approximately 10-fold higher in the brain and only 2.5-fold higher in the lungs compared to WT mice (Figure 5A), these findings suggest that distinct ISGs may contribute differentially to bacterial control in an organ specific manner. Next, we used flow cytometry to profile innate and adaptive immune cell populations in the spleens. We observed an imbalance between effector and regulatory T cells in humanized mice, characterized by an increase in activated effector T cells and a reduction in Treg cells (Figure 6). This imbalance might be due to the impaired bacterial clearance and may lead to heightened systemic inflammation. It is known that IFN signaling is critical for antigen-presenting cell maturation and NK cell activation(77–80). Our data showed that humanized IFN-γ signaling led to a significant reduction in MHC class II expression on antigen-presenting cells and impaired activation of NK cells (Figure 7). This finding reveals a potential underlying mechanism for the greater susceptibility of humans to *Ot* infection compared to mice. Neutrophilia is a common feature of acute scrub typhus(44) and increased neutrophil activation has been associated with disease progression(45). Consistent with these clinical observations, we found that neutrophil populations were more than 3-fold higher in humanized mice compared to WT mice following infection (Figure 7), suggesting that this humanized mouse model recapitulates key aspects of immune dysregulation observed in patients with acute scrub typhus. Lastly, we measured the *Ot*-specific antibody titers in mouse serum and found that humanized mice produced IgM and IgG comparable to those of WT controls (S3 Fig). The intact antibody response supports the suitability of this animal model for future vaccine studies.

Humanized mouse models represent promising tools for studying scrub typhus. Jiang et al. found that humanized DRAGA mice were highly susceptible to intradermal and subcutaneous infection, mounted strong CD4^+^ and cytotoxic T cell responses, and produced human IgM and IgG following repeated immunization, suggesting it as a new pre-clinical model for *Ot* pathogenesis study and vaccine candidate testing(50). Our study using mice with humanized IFN-γ signaling builds on our recent findings that IFN-γ, rather than type I interferons, plays a critical role in skin eschar formation and host defense against *Ot* infection(25). To our knowledge, this triple KO/KI mouse model has never been applied in any research study so far. Although the vendor validated the genetic manipulation, downstream signaling activation, and immune cell profiles, we further confirmed the expression of IFN-γ and its receptors in both humanized and WT mice during infection (Figure 3). A limitation of our study is the limited number of animals available, which restricted our ability to comprehensively assess the temporal dynamics of susceptibility, pathology, and immune responses. Since our model is a genetically engineered humanized mouse strain, it should not be confused with cellular humanization, where mice are engrafted with human cells such as hematopoietic stem cells or peripheral blood mononuclear cells. Moreover, these germline-modified mice can be bred, making them a more cost-effective and time-efficient option for experimental studies.

In sum, we reported a novel humanized mouse model that is susceptible to *Ot* infection and forms skin eschars via an intradermal infection route, effectively replicating human scrub typhus disease. Future investigations are warranted by using this novel model to evaluate bacterial virulence, characterize host protective immunity and assess the efficacy of potential new vaccine candidates.

## Acknowledgements

We would like to thank the UTMB Flow Cytometry and Cell Sorting Core Lab (Meredith Weglarz) and Dr. Joseph Jelinski for sample analysis and manuscript revision, respectively.

---

**S1 Table. Sequences of PCR primers.** All primer sequences are obtained from PrimerBank (https://pga.mgh.harvard.edu/primerbank/) or our previous publications.

**S2 Fig. Skin histology of uninfected mice**. Skin tissues from uninfected WT and humanized mice were collected and subjected to H&E staining. Representative histological images are shown at 14 dpi. Scale bar = 100 µm.

**S3 Fig. Comparable *Ot*-specific IgM and IgG levels in humanized and WT control mice.** To measure relative amounts of IgM and IgG antibodies, mouse sera were collected at 14 dpi as shown in Figure 1 and then diluted to create a dilution curve. Uninfected mouse sera were used as controls. The area under the curve (AUC) is calculated from three pooled independent experiments and shown as mean ± SD. One-way ANOVA with a Šídák’s multiple comparisons was used for data analysis. *, *p* < 0.05; **, *p* < 0.01.

**S4 Fig. Reduced expression of ISGs in the lungs of humanized mice with *Ot* infection.** Mice were infected as described in Figure 1 and were euthanized at 14 dpi. The transcript levels of ISGs and inflammatory genes in the lungs were analyzed by qRT-PCR. Data are shown as mean ± SD from three pooled independent experiments and analyzed by one-way ANOVA with a Šídák’s multiple comparisons test. *, *p* < 0.05; **, *p* < 0.01; ***, *p* < 0.001.

