## Supplementary Table 1 for "A Humanized IFN-γ Mouse Model Reveals Skin Eschar Formation, Enhanced Susceptibility and Scrub Typhus Pathogenesis"

**Real-time PCR primers of murine genes**

Forward (5’ to 3’)	Reverse (5’ to 3’)

*Ifngr1*	CTGGCAGGATGATTCTGCTGG	GCATACGACAGGGTTCAAGTTAT

*Ifngr2*	TCCTCGCCAGACTCGTTTTC	GTCTTGGGTCATTGCTGGAAG

*Cxcl10*	CCAAGTGCTGCCGTCATTTTC	GGCTCGCAGGGATGATTTCAA

*Nos2*	GTTCTCAGCCCAACAATACAAGA	GTGGACGGGTCGATGTCAC

*Ifit1*	CTGAGATGTCACTTCACATGGAA	GTGCATCCCCAATGGGTTCT

*Ifit2*	AGTACAACGAGTAAGGAGTCACT	AGGCCAGTATGTTGCACATGG

*Gbp2*	CTGCACTATGTGACGGAGCTA	GAGTCCACACAAAGGTTGGAAA

*Gbp4*	GGAGAAGCTAACGAAGGAACAA	TTCCACAAGGGAATCACCATTTT

*Gbp5*	CAGACCTATTTGAACGCCAAAGA	TGCCTTGATTCTATCAGCCTCT

*Isg15*	GGTGTCCGTGACTAACTCCAT	TGGAAAGGGTAAGACCGTCCT

*Isg20*	TGGGCCTCAAAGGGTGAGT	CGGGTCGGATGTACTTGTCATA

*Stat1*	TCACAGTGGTTCGAGCTTCAG	CGAGACATCATAGGCAGCGTG

*Rsad2*	TGCTGGCTGAGAATAGCATTAGG	GCTGAGTGCTGTTCCCATCT

*Oas2*	TTGAAGAGGAATACATGCGGAAG	GGGTCTGCATTACTGGCACTT

*Oas3*	TCTGGGGTCGCTAAACATCAC	GATGACGAGTTCGACATCGGT

*Oas1b*	GGGCCTCTAAAGGGGTCAAG	TCAAACTTCACTCCACAACGTC

*Oasl2*	TTGTGCGGAGGATCAGGTACT	TGATGGTGTCGCAGTCTTTGA

*Il6*	TAGTCCTTCCTACCCCAATTTCC	TTGGTCCTTAGCCACTCCTTC

*Il27*	CTGTTGCTGCTACCCTTGCTT	CACTCCTGGCAATCGAGATTC

*Tnf*	CCCTCACACTCAGATCATCTTCT	GCTACGACGTGGGCTACAG

*Sele*	ATGCCTCGCGCTTTCTCTC	GTAGTCCCGCTGACAGTATGC

*Selp*	CATCTGGTTCAGTGCTTTGATCT	ACCCGTGAGTTATTCCATGAGT

*Ifitm3*	CCCCCAAACTACGAAAGAATCA	ACCATCTTCCGATCCCTAGAC

*Icam1*	GTGATGCTCAGGTATCCATCCA	CACAGTTCTCAAAGCACAGCG

*Vcam1*	AGTTGGGGATTCGGTTGTTCT	CCCCTCATTCCTTACCACCC

*Endothelin-1*	GCACCGGAGCTGAGAATGG	GTGGCAGAAGTAGACACACTC

*Vegf*	CTGCCGTCCGATTGAGACC	CCCCTCCTTGTACCACTGTC

*Gapdh*	TGGAAAGCTGTGGCGTGAT	TGCTTCACCACCTTCTTGAT

**Real-time PCR primers of human genes**

Forward (5’ to 3’)	Reverse (5’ to 3’)

*IFNGR1*	TCTTTGGGTCAGAGTTAAAGCCA	TTCCATCTCGGCATACAGCAA

*IFNGR2*	CTCCTCAGCACCCGAAGATTC	GCCGTGAACCATTTACTGTCG
