## Supplementary figures and images for "A Humanized IFN-γ Mouse Model Reveals Skin Eschar Formation, Enhanced Susceptibility and Scrub Typhus Pathogenesis"

### Supplementary Figure 2

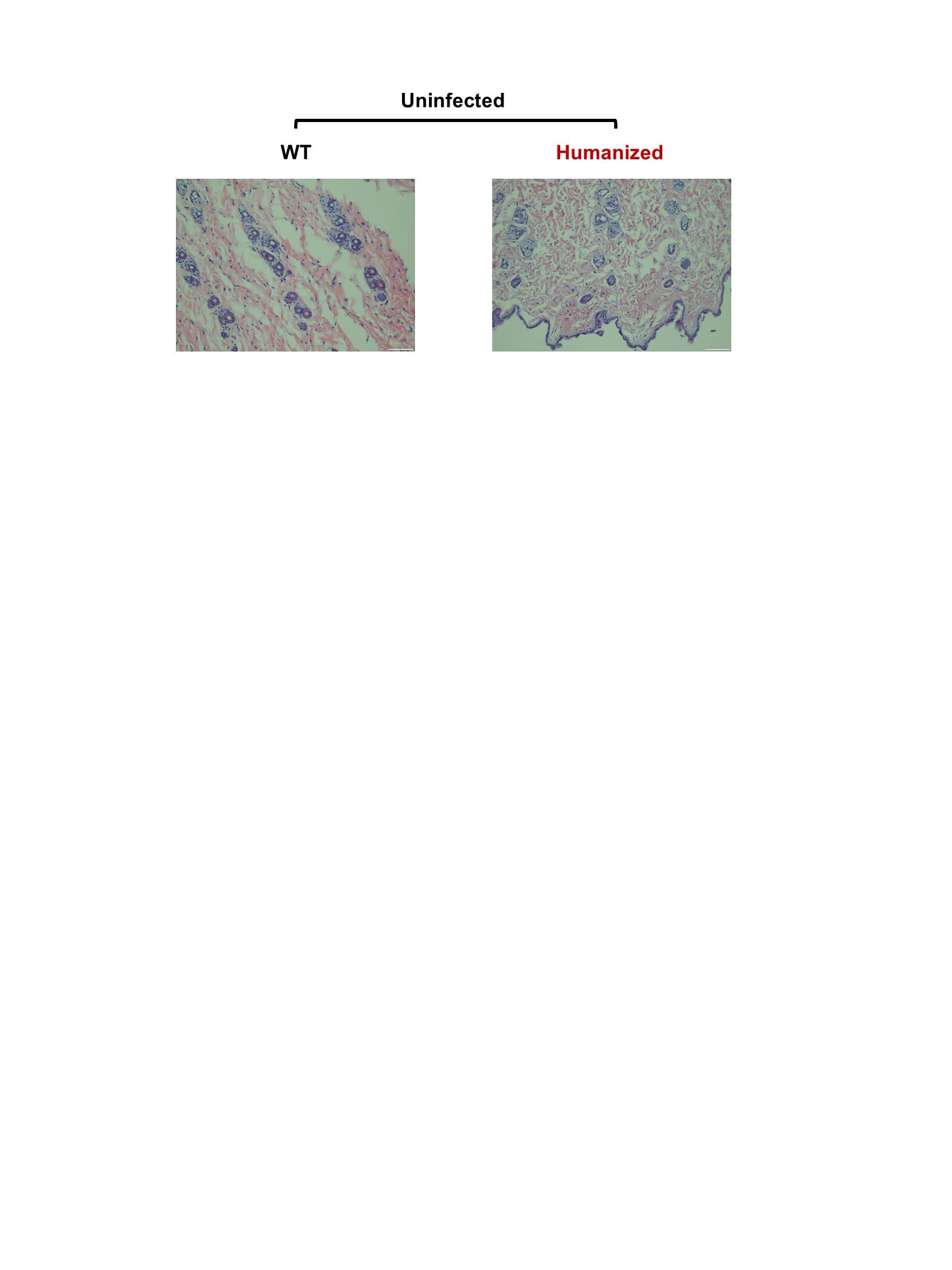

### Supplementary Figure 3

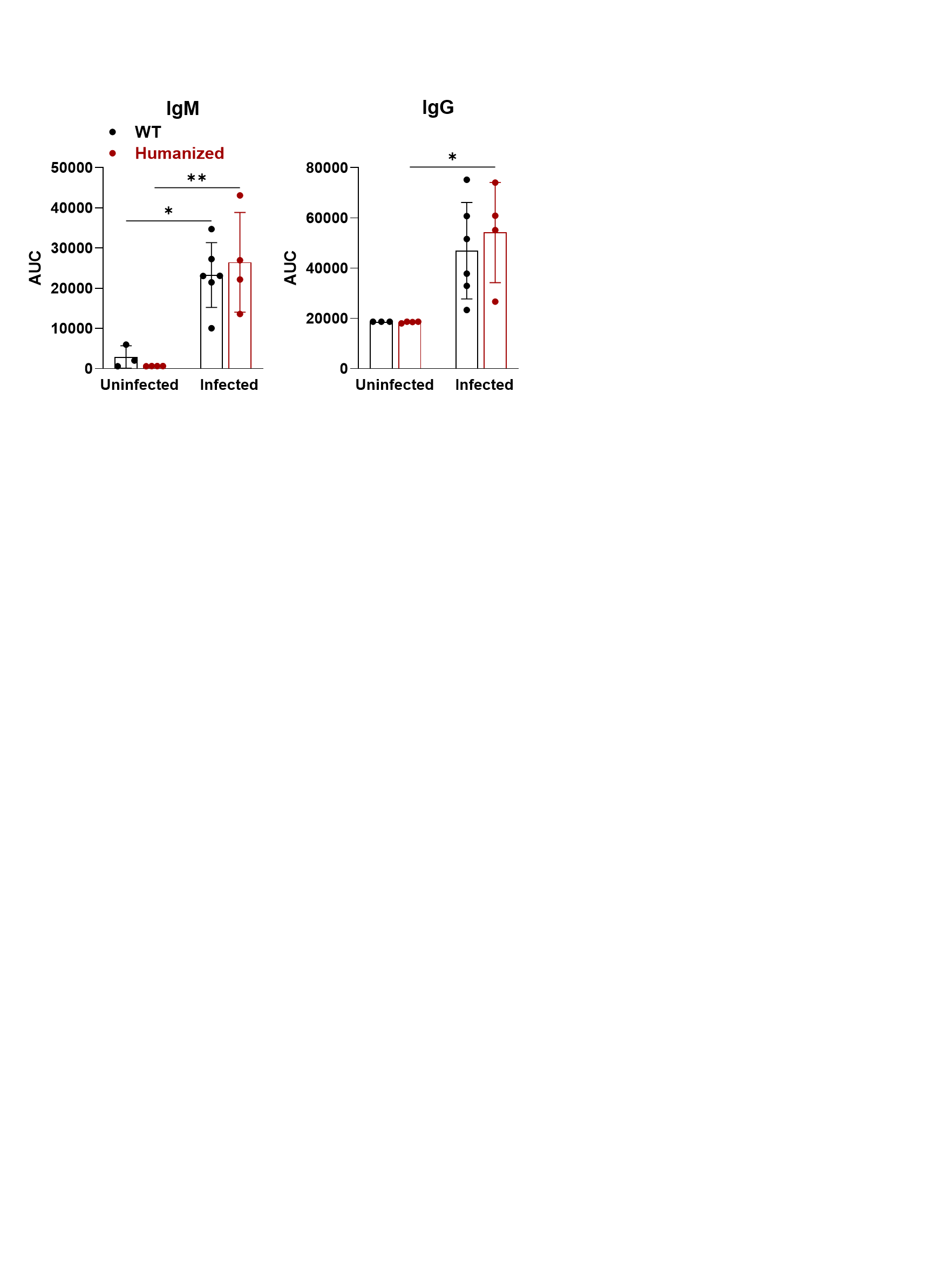

### Supplementary Figure 4

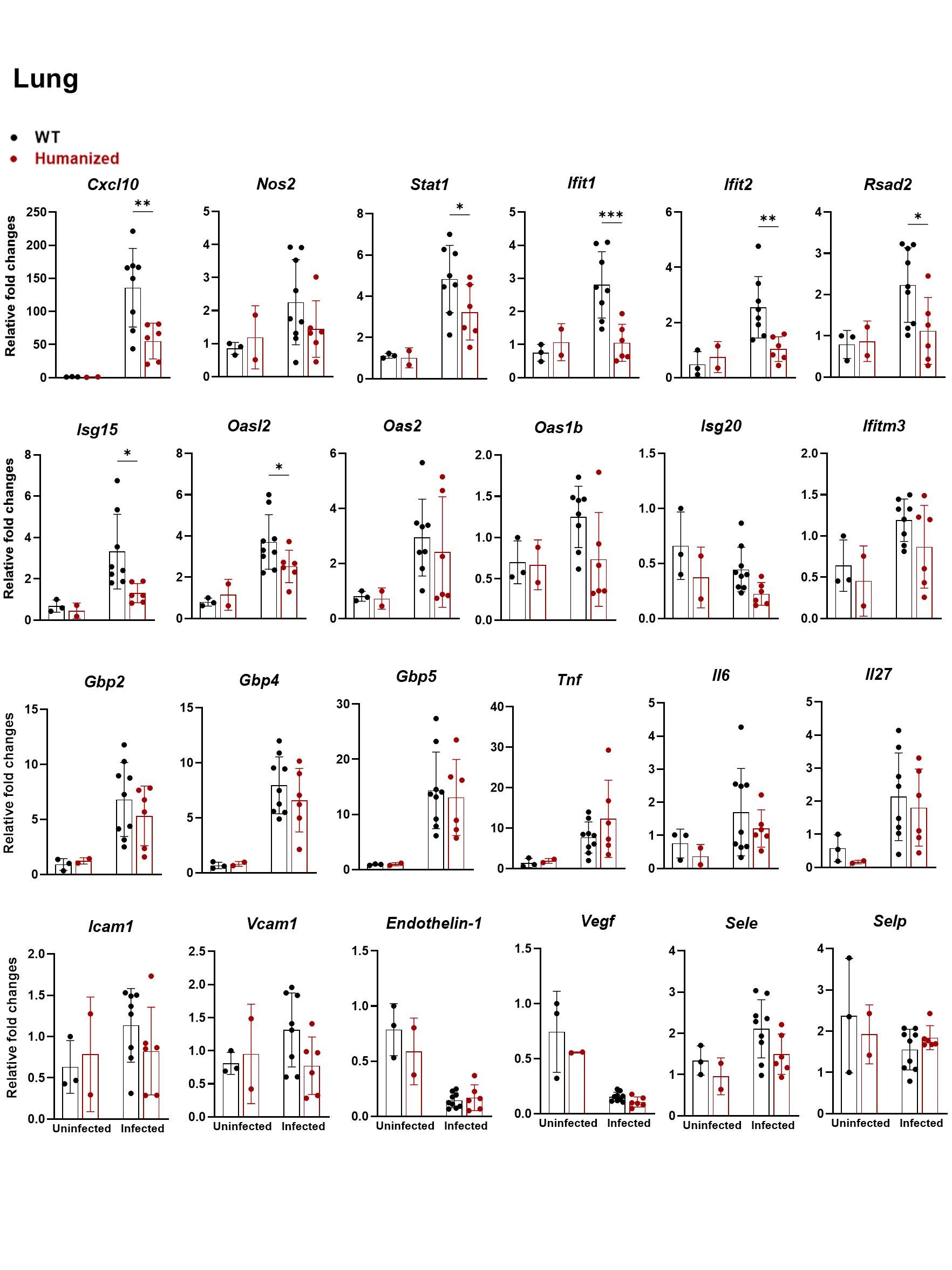
